# Riparian forests shape trophic interactions in detrital stream food webs

**DOI:** 10.1101/2023.10.31.564911

**Authors:** Rebecca Oester, Florian Altermatt, Andreas Bruder

## Abstract

Freshwater and terrestrial biodiversity is linked through resource flows. For example, subsidies from the riparian vegetation form the base of food webs in small streams. Despite the key role of detritivores in these food webs, consequences of altered resource availability and riparian vegetation type on their trophic strategies are largely unknown. Therefore, we experimentally tested direct and indirect effects of riparian vegetation type on trophic interactions and dietary imbalances of detritivores. We used stoichiometric and isotopic differences between consumers and resources as functional measures of trophic link strength. Our results show that the lack – compared to the presence – of riparian forests directly affected both stoichiometric and isotopic differences in detrital food webs, yet with diverging patterns between resources and consumers, ultimately leading to aquatic-terrestrial decoupling. Consequently, our findings demonstrate that riparian forests are essential for aquatic food webs by influencing both organisms and interactions networks.

## Introduction

While global biodiversity is declining at an unprecedented rate in both aquatic and terrestrial ecosystems (IPBES 2019; Pereira *et al*. 2010), transition zones between adjacent ecosystems are still local hotspots and refuges for biodiversity (Meyer *et al*. 2007; Risser 1995; Smith *et al*. 1997) through subsidy flows (Baxter *et al*. 2005; Scherer-Lorenzen *et al*. 2022). High rates of cross-ecosystem fluxes of nutrients and energy can create tight linkages and promote transboundary reliance of organisms and processes (Barnes *et al*. 2018). These cross-ecosystem linkages are universal (Gounand *et al*. 2018), yet, they are threatened by anthropogenic pressures (Doi 2009; Larsen *et al*. 2016). For example, land-use change affecting riparian forests not only influences terrestrial biodiversity, but may also cascade to bordering aquatic ecosystems through changes in microclimate and subsidy flows (Baxter *et al*. 2005; Webster & Meyer 1997). Deforestation of riparian forests, therefore, has potentially large effects on aquatic and terrestrial biodiversity across trophic levels (Kaylor & Warren 2017; Wallace *et al*. 1997) and ecosystem functioning (England & Rosemond 2004; Little & Altermatt 2018; Warren *et al*. 2016). In particular, these effects are detrimental to species and processes directly relying on riparian forests (England & Rosemond 2004; Kominoski *et al*. 2011; Little & Altermatt 2018).

The base of many stream food webs is largely subsidized by riparian vegetation through input of leaf litter and other organic matter (Collins *et al*. 2016; Marcarelli *et al*. 2011). Despite leaf litter being relatively nutrient poor (Leberfinger *et al*. 2011; Ferreira *et al*. 2016; Brett *et al*. 2017), terrestrial inputs are essential for these typically heterotrophic ecosystems (Rosi-Marshall & Wallace 2002; Vannote *et al*. 1980; Webster & Meyer 1997). There, a suite of microbial decomposers and detritivores process vast quantities of allochthonous plant material throughout the year (Boyero *et al*. 2021; Gessner *et al*. 2010). When leaf litter falls into streams, the first to colonize these resources are microorganisms, in particular fungi (Findlay & Arsuffi 1989; Hieber & Gessner 2002). Microbial decomposers subsequently consume the leaf litter and make it also more palatable to other consumers (Brosed *et al*. 2017; Danger *et al*. 2012; Fuller *et al*. 2015), by immobilizing and accumulating nutrients (Arsuffi & Suberkropp 1989; Cross *et al*. 2003). Many detritivorous consumers, such as the functional feeding group of macroinvertebrate shredders (*sensu* Moog, 1995), feed on this plant litter conditioned by microbes (Arsuffi & Suberkropp 1989; Cummins *et al*. 1989). This shredding feeding mode evolved across many taxa of aquatic insects (e.g., Plecoptera or Trichoptera) and other macroinvertebrate orders (e.g., Amphipoda). Eventually, this detrital food web, containing dead organic matter, microbial decomposers and detritivores, supports and connects higher trophic levels in the aquatic and terrestrial ecosystem, such as fish or birds (Allen *et al*. 2012; Twining *et al*. 2019; Woodward *et al*. 2008).

Theory in ecological stoichiometry predicts that, when a consumer’s diet does not meet its physiological needs, consumer fitness, growth or behaviour is affected (Cruz-Rivera & Hay 2003; Sterner & Elser 2002; Vanni 2002). While many consumers can be in nutritional imbalance with their resources temporarily (Lemoine *et al*. 2014), detritivores often show a large dietary imbalance with their dominant source of food, i.e. leaves and wood (Atkinson *et al*. 2022). Despite behavioral strategies like selective or compensatory feeding and metabolic adaptations, imbalances between consumers and their food often remain (Frainer *et al*. 2016). Limitations of certain elements or specific compounds and spatio-temporal asynchrony of resource availability can influence different dimensions of this imbalance (Frainer *et al*. 2016; Labed-Veydert *et al*. 2022; Larsen *et al*. 2016).Yet, the degree of dietary imbalance and environmental factors influencing these differences for different detritivore taxa are largely unknown (but see, Burdon *et al*. 2020; Frainer *et al*. 2016; Lemoine *et al*. 2014). Moreover, trophic interactions and dietary strategies for different taxa in ecosystems at the land-water interface are not well understood *in situ*. Incorporating trophic interactions in biodiversity studies (Duffy *et al*. 2007; Jacquet *et al*. 2022) can improve understanding of aquatic-terrestrial linkages (England & Rosemond 2004), cross-ecosystem nutrient flows (Gounand *et al*. 2018) and their importance for key consumers and ecosystem processes (Frainer *et al*. 2016; Little *et al*. 2019).

We studied trophic interactions of detritus-based communities in streams and disentangled the direct and indirect effects of riparian vegetation type on resources and consumers, their stoichiometric and isotopic composition, and dietary imbalances. To do so, we exposed leaf litter in different headwater streams with forested or non-forested riparian vegetation to be naturally colonized and consumed by microbial decomposers and detritivores. We measured Carbon (C) to Nitrogen (N) ratios to assess organismal tissue composition (Cross *et al*. 2003; Sterner & Elser 2002) in addition to *δ*13C signatures to trace the source of carbon through the food web (Boecklen *et al*. 2011). We used stoichiometric and isotopic differences between resources and invertebrates as two dimensions of dietary imbalances and as a functional measure of trophic links. Since dietary imbalance of consumer individuals might be linked allometrically to metabolism, nutrient cycling and mode of resource acquisition (García *et al*. 2017; García‒Girón *et al*. 2022; Krynak & Yates 2018) we additionally measured the mass of shredders and aquatic fungi. We expected direct effects of riparian vegetation type (forested vs. non-forested) on (1) consumer mass (fungal biomass and mean shredder body mass), and food web (2) stoichiometry and (3) isotopic signatures. Moreover, we expected indirect effects of riparian vegetation type over consumer mass on (4) C:N ratios and (5) *δ*13C signatures, respectively (Figure 1, Table S1). By quantifying the direct and indirect influence of riparian vegetation type on aquatic ecosystems, we provide key findings for assessing the consequences of deforestation in riparian zones and ultimately for guiding restoration and management efforts.

**Figure 1:**
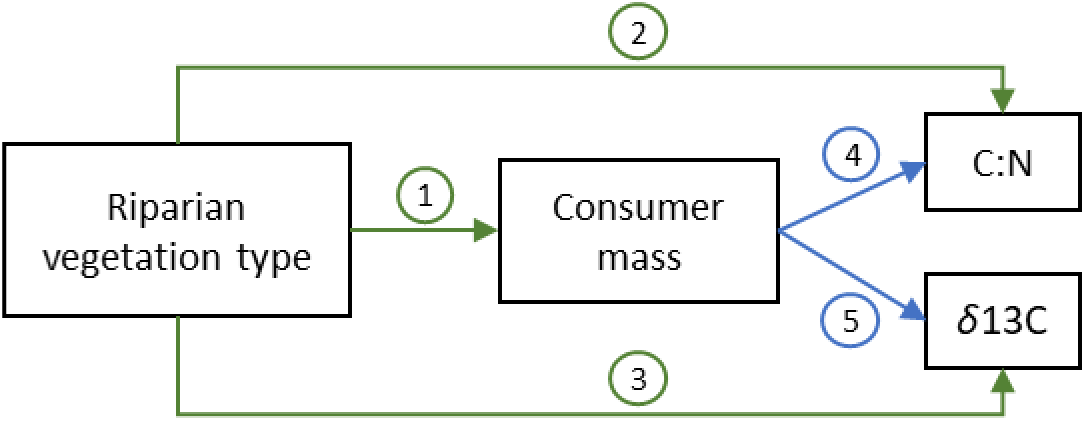
Conceptual meta-model with direct and indirect effects (arrows) of riparian vegetation type (forested vs. non-forested) and consumer mass on C:N ratios and *δ*13C values of resources and consumers (boxes). The encircled numbers 1–5 represent hypothesized paths explained in detail in Table S1. The green paths 1, 2 and 3 represent direct effects from riparian vegetation type to consumer mass, C:N ratios and *δ*13C signatures, respectively. The blue paths 4 and 5 show indirect paths from riparian vegetation type over consumer mass to stoichiometry and stable isotope signature, respectively.

## Material and Methods

### Study sites

We studied detritus-based communities and their trophic interactions in eight 1^st^ to 3^rd^ order headwater streams in Switzerland in autumn/winter 2020/2021. We selected eight streams with a forested section (mixed deciduous trees) and a non-forested section (grassland and/or extensively used pasture) in a paired design of approximately 500 m apart (Oester et al. 2023). The herein-studied contrast of forested vs. non-forested stream sections is a commonly found pattern at landscape scales, with well-documented effects on the composition of aquatic detritivore communities, and representative for the studied stream systems (Cereghetti & Altermatt 2023; Little & Altermatt 2018). The two sites and tributaries of each of the eight streams showed no to minor anthropogenic impacts and did not substantially differ in average water temperature (forested: 4.2 °C ± 1.1; non-forested: 4.4 °C ± 1.4), nitrate concentrations (forested: 2.4 mg/l ± 2.1; non-forested: 2.2 mg/l ± 2.1), or DOC (forested: 2.7 mg/l ±1.9; non-forested: 2.7 mg/l ± 2.0) (for more details on the site characteristics and invertebrate communities see Table S2 and Oester *et al*. (2023)).

### Data collection

To examine detritus-based communities in streams, we deployed 12 leaf litter bags per site as subsidized resources. These bags consisted of coarse mesh (1 cm) allowing microbial decomposers and macroinvertebrates to colonize and consume the leaves. We used 5 g (± 0.03 g) of dried leaf litter (dried at 40 °C for 48 h) of two common riparian tree species: black alder (*Alnus glutinosa* (L.) Gaertn.) and European ash (*Fraxinus excelsior* L.), and an (equal mass) mixture of both leaf species to represent three resource options (Bruder *et al*. 2014). As there were no statistically significant differences in shredder abundance, average dry weight, or total biomass on the three different resources, we henceforth grouped resources only by leaf species and averaged the values of the mixed leaf litter treatment for subsequent analyses.

To assess microbial colonization and its effects on resource quality, we quantified fungal biomass in all leaf litter bags (Gessner 2020). First, we extracted ergosterol (as a proxy for fungal biomass) from freeze-dried leaf disks from the experimental leaves. We subsequently purified these extracts with solid-phase extraction (Sep-Pak® Vac RC tC18 500 mg sorbent; Waters, Milford, USA). We measured the ergosterol concentration using ultra-high-performance liquid chromatography (UHPLC; 1250 Infinity Series, Agilent Technologies, Santa Clara, USA) at a wavelength of 282 nm and calculated fungal biomass with a conversion factor of 5.5 mg ergosterol per g fungal biomass according to Gessner (2020).

To characterize leaf-associated detritivore assemblages, we collected all macroinvertebrates inside each leaf litter bag at the timepoint when on average ∼50 % of all the leaf litter was decomposed (i.e., after approximately one month). We excluded data from leaf litter bags with < 25 % leaf litter mass remaining, as retention of macroinvertebrates in bags with limited leaf litter is minimal. We preserved the macroinvertebrates in ethanol (98 %) and identified all individuals under a dissecting microscope to the lowest taxonomic level possible (i.e., mostly genus or subfamily; e.g., Sundermann et al., 2007; Tachet et al., 2010). For each taxon, we assigned feeding preference (Schmidt-Kloiber & Hering, 2015; version 8.0; accessed on 01 October 2021). We focused on the functional feeding group of shredders, whose preferred resource is coarse particulate organic matter, i.e., leaf litter and other detritus *sensu* Moog (1995). Shredders from three major taxa, i.e., Crustacea, Plecoptera and Trichoptera were most common and abundant, and included *Gammarus fossarum* (Crustacea)*, Capnia* sp., *Leuctra* sp., *Nemoura* sp., *Protonemura* sp. (Plecoptera), Limnephilini and Limnephilidae group Auricollis (Trichoptera). We measured body length to estimate macroinvertebrate dry weight using taxon-specific length-weight regression models (e.g., Benke et al., 1999; Baumgärtner & Rothhaupt, 2003). We averaged the dry weight of all the individuals of each order in each leaf litter bag to incorporate body mass as an additional variable in subsequent analyses. Together these seven taxa contributed 63 % of the total invertebrate biomass in the leaf litter bags. In 70 % of all leaf litter bags there was sufficient invertebrate biomass of at least one taxon to analyze trophic interactions using stoichiometric and stable isotope analyses.

### Stoichiometry and stable isotope analysis

To characterize trophic interactions between resources and consumers, we analyzed the stable isotopes and total contents of carbon and nitrogen with an Elemental Analyzer and Isotope Ratio Mass Spectrometer (EA-IRMS; Elementar, Vario Pyro; IsoPrime). For each bag, we dried (60 °C for 48 h), homogenized, and encapsulated (9 × 5-mm tin capsules, Säntis Analytical) the tissues of leaves and shredders. To assess the leaf composition at the initial stage (T0), we measured a subset of leaf material for each leaf species from the same batch as that used for the field experiment. For the measurements to lie in the detection range of the EA-IRMS, we used 1–1.5 mg of leaves, and 0.8–1.5 mg of macroinvertebrate tissue. To fall within these weight limits, we aggregated 3–100 whole shredder individuals per taxon per leaf litter bag, depending on abundances and estimated dry weight. We report the results as *δ*13C defined as the ‰ deviation of the sample from a reference (i.e., Pee Dee belemnite carbonate) and the C:N ratios in log-transformed molar proportions (Isles 2020). For all aquatic invertebrates, we used an enrichment factor of 0.1 (Brauns *et al*. 2018) (detailed methods for *δ*13C in Supplementary Material; Figure S3).

### Quantifying dietary imbalance

To assess two important dimensions of dietary imbalance between resources and consumers, we used two complementary metrics: the difference in *δ*13C signatures and the difference in C:N ratios between shredder body tissue and the resource (i.e., experimental leaf litter conditioned by microbes). To quantify isotopic difference, we subtracted the *δ*13C values of leaf litter from *δ*13C values of shredders (Burdon et al., 2020). We tested different approaches to calculate C:N differences, including subtractions (e.g., Cross et al., 2003; Hladyz et al., 2009) and LRR (e.g., Frainer et al., 2016), and ratios (e.g., Mooshammer et al., 2014) between shredders and leaf litter, which led to very similar results (Figure S2). We here report C:N differences as ratios between resources and consumers. The higher these isotopic and stoichiometric differences, the higher the dietary imbalance between resource and consumer.

### Statistical analysis

We performed all calculations and statistical analyses in R (version 4.1.2; R Development Core Team, 2020). To assess the relative importance of direct and indirect links between riparian vegetation type (levels: forested and non-forested) on C:N ratios and *δ*13C signatures of resources and consumers in the detrital food web, we constructed Bayesian generalized non-linear multivariate multilevel models (brms; Bückner 2019) and visualized the relationships through path diagrams. We included indirect effects of riparian vegetation type over consumer mass based on our hypotheses described in Table S1. We used the same model structure for all resources and consumers (Figure 1) for consistent comparisons. We scaled all numeric values (mean = 0) and included weakly informative priors of normally distributed intercepts (0, 1) and slopes (0, 0.1). To account for the study design and environmental differences between streams, we included sampling location as random effect. For each resource and consumer group, we then tested the direct effects on C:N ratios and *δ*13C signatures and compared the stoichiometric and isotopic difference between forested and non-forested sites using the same model specifications as described above. For all models, we generated 20,000 (four chains run for 10,000 iterations discarding the first 5,000 as burn-in) Markov chain Monte Carlo (MCMC) samples from the posterior distribution where draws were sampled using NUTS (No-U-Turn Sampler). All MCMC chains indicated convergence through falling within the threshold specified by Gelman and Rubin (1992) and showing high effective sample size measures. We visually checked the fit of the posterior distribution with the data (bayesplot, Gabry et al., 2021).

## Results

We found strong evidence for direct effects of riparian vegetation type on stoichiometry and isotopic signatures of key resources and consumers of stream food webs (Figure 2). The direction and strength of these direct effects, however, differed between resources and different consumer groups. In the absence of riparian forests, resources (i.e., alder and ash leaf litter colonized by microbes) and Trichoptera shredders showed significantly lower C:N ratios (Figure 2,3) due to higher %N in these tissues (Figure S1B), while Crustacea and Plecoptera stoichiometry was unaffected by riparian vegetation type. Contrastingly, Crustacea and Plecoptera shredders from non-forested sites showed lower *δ*13C signatures compared to individuals from forested sites (Figures 2,3). Microbial conditioning, measured as fungal biomass, was consistently higher in the non-forested sites compared to the paired forested sites (Alder: estimate = 0.16 [0.06, 0.25]; Ash: estimate = 0.11 [0.05, 0.22]). However, we found no evidence for any indirect effects from riparian forests over consumer mass to either stoichiometry or isotopic signatures (Figure 2).

**Figure 2:**
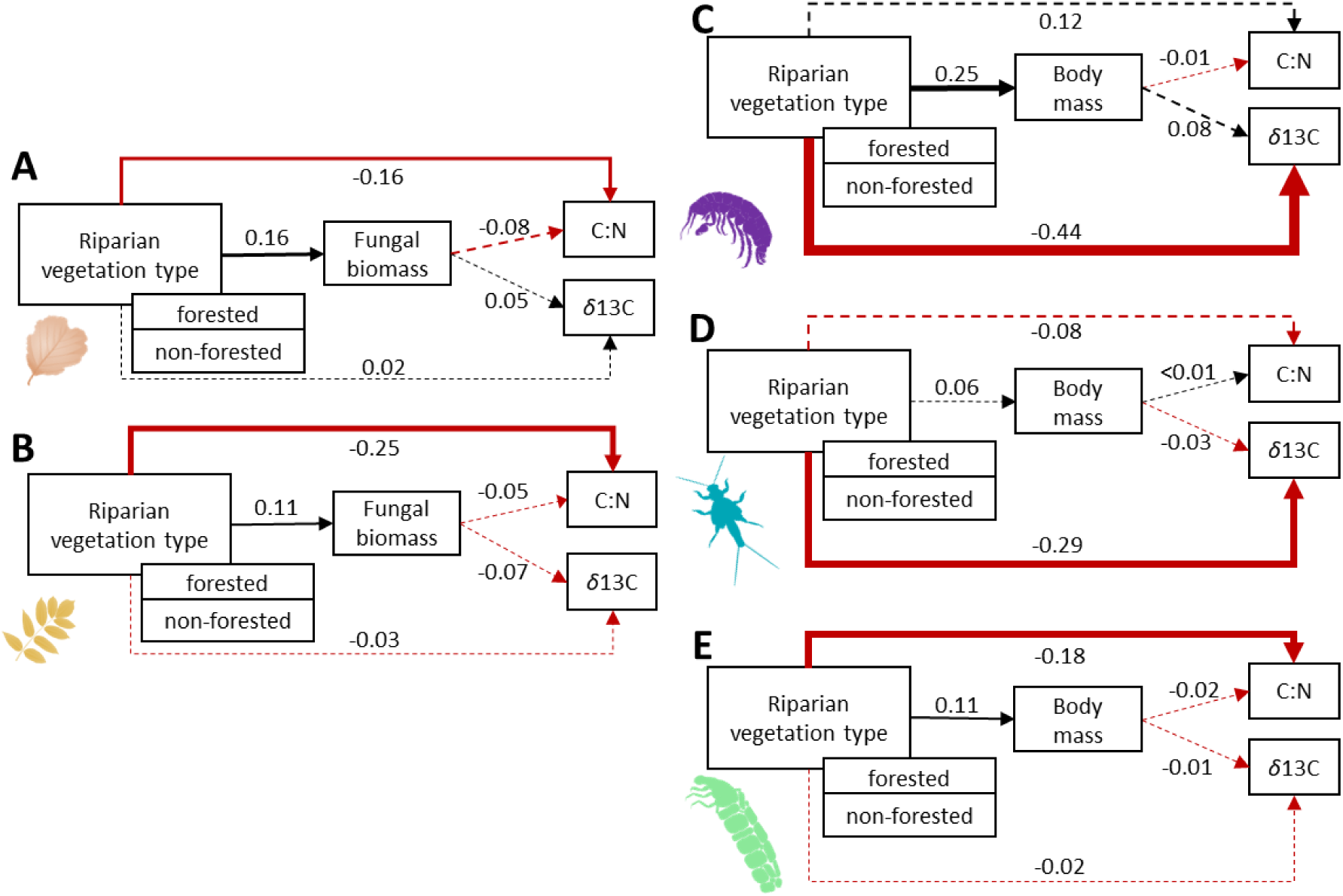
Path analysis of direct and indirect (over consumer mass) effects of riparian vegetation types: A: fungal colonized alder, B: fungal colonized ash, C: Crustacean shredders, D: Plecopteran shredders, and E: Trichopteran shredders. Solid paths indicate strong evidence for these relationships (i.e., 0 not included in CIs), while dashed paths represent no evidence for effects. Red paths indicate negative relationships while black paths indicate positive effects. Arrows from riparian vegetation type refer to differences from forested (reference) to non-forested. The values for each path refer to scaled values (i.e., the higher the value, the stronger the relative effect strength).

**Figure 3:**
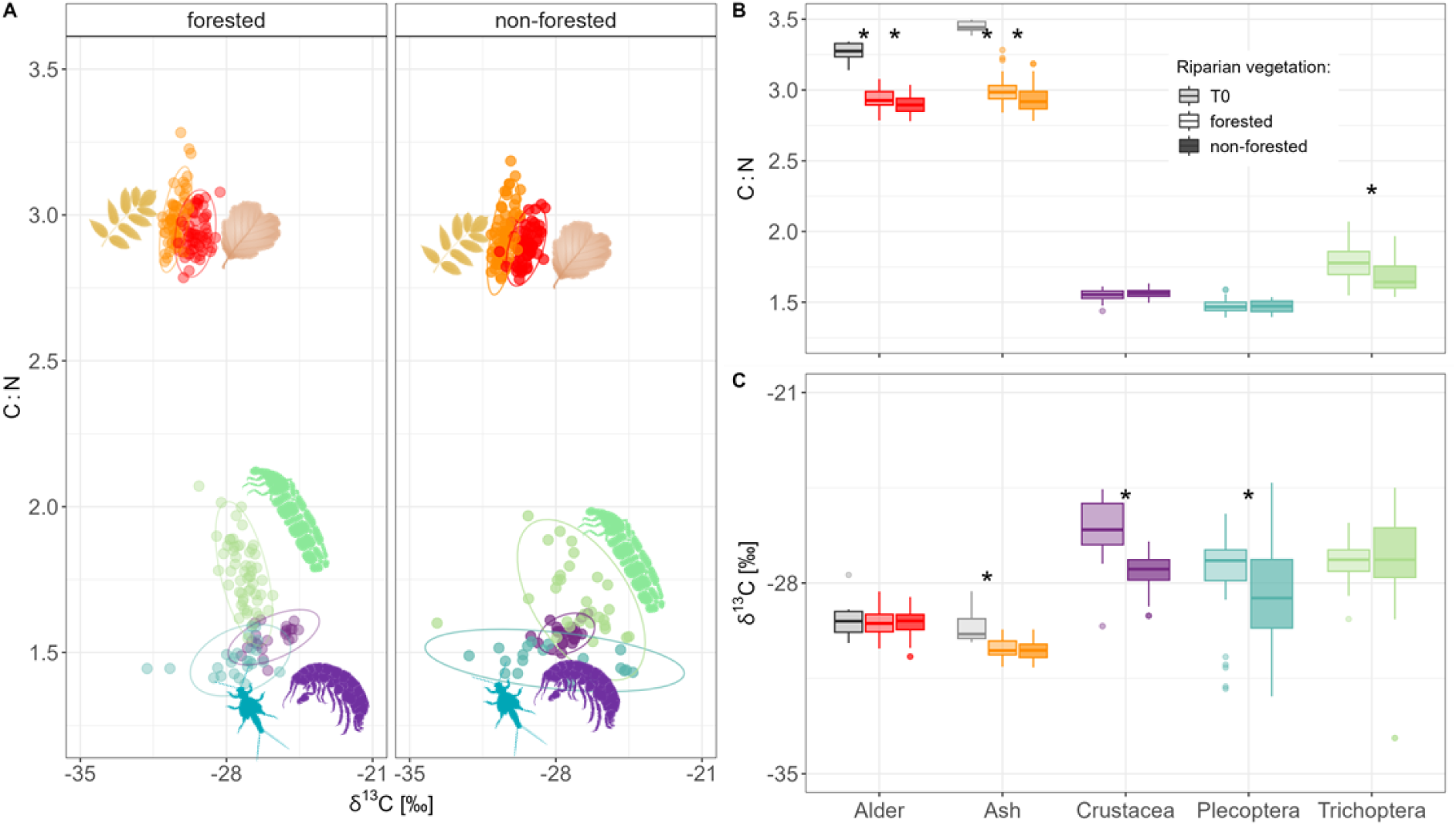
Total effects of riparian vegetation type on A) detrital community C:N ratios (ln molar ratios) and δ13C signatures between forested and non-forested sites (ellipses show the 95% confidence area), B) C:N ratios (ln molar ratios) and C) δ13C signatures of alder, ash, Crustacea, Plecoptera and Trichoptera. Asterisks indicate statistically significant differences (i.e., 0 not included in CIs).

C:N ratios of leaves differed between before and after the experiment, and between forested and non-forested sites revealing the effects of conditioning and microbial processing (Figure 3). From the initial 1.6 %N (ash) and 2.2 %N (alder), both leaf species increased 1%N during the field experiment (Figure S1B). Additionally, alder lost 2 % from the initial 48.6 %C (Figure S1A). Consequently, after stream exposure, C:N ratios of leaf litter were lower than at start and became more similar between the leaf species (Figure 3A,B). Compared to the C:N ratios of the initial leaf litter, the shredder C:N values were approximately 2.1 – 2.2 times lower. However, the difference in C:N ratios between conditioned leaf litter and shredders was only 1.3 – 1.8 times lower due to higher %N accumulated in the leaves (Figure 3B). Trichoptera shredders showed the highest mean logC:N ratio (1.73 ± 0.12) compared to Crustacea (1.56 ± 04) and Plecoptera (1.47 ± 0.05).

Before exposure, *δ*13C values of alder and ash litter showed substantial overlap (Alder: −29.34 ± 0.71; Ash: −29.56 ± 0.61; Figure 3A,C). While *δ*13C signatures of alder did not change during the experiment, *δ*13C values of ash shifted on average −0.85 ‰. Consequently, after the field experiment, alder showed 1 ‰ higher *δ*13C values than ash. However, there was no effect of riparian vegetation type on *δ*13C values within each leaf treatment (Figure 3C). Overall, the three shredder orders showed great overlap in the mean carbon signatures (Crustacea: −26.89 ‰ ± 1.22; Plecoptera: −28.12 ‰ ± 1.66; Trichoptera: - 27.23 ‰ ± 1.09).

For all shredders, we found substantial stoichiometric imbalances between their body tissue and resources with approximately double the C:N values of body tissue compared to that of leaves (Figure 3B, 4A-D). Resulting from differing *δ*13C and C:N patterns between resources and consumers, the dietary imbalances were most different between forested and non-forested sites in Crustacea (Figure 4A,E). Crustacea showed both 45 % and 4 % bigger differences in *δ*13C signatures and C:N ratios, respectively, between forested and to non-forested sites. Plecoptera shredders showed 4 % higher *δ*13C differences (Figure 4E). For Trichoptera, we found small trends for the opposite pattern with, for example, 3 % lower C:N differences in the forested sites (Figure 4A).

**Figure 4:**
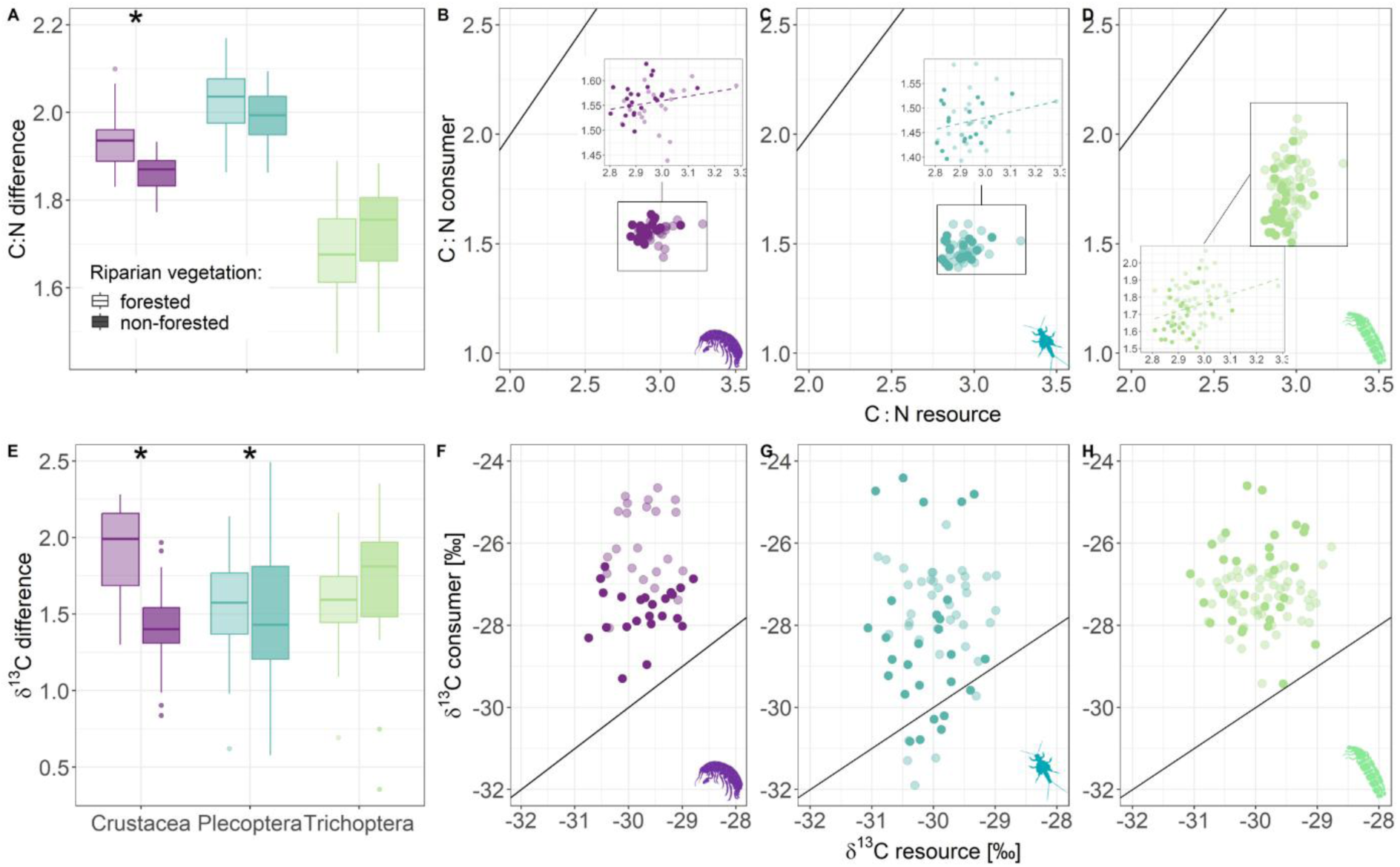
Dietary imbalances expressed as stoichiometric (A-D) and isotopic (E-H) differences between resources and consumers. A) and E) show the effects of riparian vegetation type on stoichiometric and isotopic differences, respectively. Asterisks indicate statistically significant differences of riparian vegetation type (i.e., 0 not included in CIs). B), C) and D) show degree of homeostatic relationships between resource C:N and consumer C:N values for Crustacea, Plecoptera and Trichoptera (as ln molar ratios), respectively. F), G) and H) show relationships between resource and consumer *δ*13C signatures. The black line indicates the 1:1 line where resource and consumer values would be equal and deviations from this line indicate dietary imbalances.

## Discussion

We found major direct effects of riparian vegetation type on aquatic food webs when contrasting forested versus non-forested stream sections. While vegetation type had strong stoichiometric effects on the base of the food web, consumers only partially matched these patterns. Instead, the type of riparian vegetation had strong isotopic effects on these consumers, revealing differences in dietary strategies and degree of reliance of consumers on local resources. In addition to confirming the great importance of cross-ecosystem resource flows from the riparian vegetation into aquatic ecosystems, our study design allowed us to quantify the differences in dietary imbalance between key consumers. Consequently, our findings highlight how effects by riparian vegetation type around streams cascade through the detrital food web by not only influencing the individual food web components but also their interactions network.

### Consumers

Decomposers and detritivores are key consumers in small streams and connect aquatic and terrestrial ecosystems through their trophic interactions (Table S1, Figure 3). Microbial decomposers, such as aquatic fungi, showed higher biomass in non-forested sites in both leaf species (Figure 2) despite very similar average nutrient and temperature levels between forested and non-forested sites within each stream in our study (see Methods). This difference in fungal biomass could come from positive priming effects of periphyton facilitating fungal growth (Danger *et al*. 2013; Halvorson *et al*. 2019a), as background periphyton biomass was greater in non-forested sites (data not shown). Alternatively, non-forested sites could have had slightly higher nutrient levels in particulate or bound form that we did not detect in the water samples. However, independent of effects of riparian vegetation type, higher fungal biomass tended to increase %N of leaves likely through conditioning (Figure S1B, Costantini *et al*. 2014; Brosed *et al*. 2017). These conditioning effects become particularly apparent when comparing stoichiometry from freshly abscessed leaves and leaves exposed in streams (Fig. 3; Manning *et al*. 2015). Consequently, this nutrient enrichment through microbial conditioning reduced the mismatch in elemental composition between resource and consumers (Fig. 3B). As corroborated by other studies, microbial conditioning is essential not only for higher trophic levels but the entire process of organic matter decomposition (Cross *et al*. 2005; Graça 2001; Graça *et al*. 1993).

While differing stoichiometric requirements and resource fidelity have been reported for differing invertebrate feeding guilds (Cross *et al*. 2003) and in lab experiments (Halvorson *et al*. 2019b), our results show clear intraguild differences in the field (Figure 2, 3). In different developmental stages, resource switching and changes in food preferences are common in stream food webs (Dick 1995; Dobson & Hildrew 1992; Wallace *et al*. 1999) which can also be influenced by density of con-specifics (Little *et al*. 2019) and subsidy influx (Leberfinger *et al*. 2011). While shredder C:N ratios were independent of their body mass (Figure 2), other elements such as phosphorous or essential biochemicals (e.g., fatty acids) might vary more between different life stages, as these compounds might relate to growth (Cross *et al*. 2005; Sterner & Elser 2002; Twining *et al*. 2019). To better understand nutrient cycling, combining theory of ecological stoichiometry with metabolic theory has been proposed (Allen & Gillooly 2009) and only recently implemented in assessing aquatic food web dynamics (Moorthi *et al*. 2016; Welti *et al*. 2017) creating interesting new research avenues.

### Shifts in trophic interactions

Stoichiometric theory predicts that resource quantity and quality influence consumer fitness and behaviour (Cruz-Rivera & Hay 2003; Sterner & Elser 2002) but homeostatic constraints and resource availability limit consumer strategies (Figure 3). For consumers with high feeding rates, such as Trichoptera shredders (Arsuffi & Suberkropp 1989; Danger *et al*. 2012), we found lower C:N values of these consumers in the forested compared to non-forested sites, mirroring the C:N patterns observed in leaf litter. This parallel stoichiometric shift of resource and consumer together with similar *δ*13C signatures between forested and non-forested sites indicate a strong reliance of Trichoptera consumers on these resources. Contrastingly, Crustacea and Plecoptera shredders exhibited no shift in C:N ratios but a higher degree of flexibility in their *δ*13C signatures with lower *δ*13C values in non-forested sites. Interestingly, these patterns differed more between the two insect orders Plecoptera and Trichoptera compared to Crustaceans despite a much smaller phylogenetic distance. This partially diverging shift between resources and consumers in a non-forested site might ultimately result in a decoupling within the detrital food web. Studies have demonstrated that land-cover changes in the riparian zone can shift stream macroinvertebrates towards a more generalist feeding behavior (Estévez *et al*. 2020) with a greater overlap of trophic niches in the community compared to forested stream sections (Burdon *et al*. 2020; Castro *et al*. 2016) and overall smaller trophic diversity (Effert-Fanta *et al*. 2022). The consequences of such altered food-web configuration and feeding niches on, for example, the fitness and consequently evolution of the consumer populations in a larger spatial and temporal context or feeding strategies of higher trophic levels in and around streams are yet to be investigated.

### Dietary imbalances

Our results show that stoichiometric and isotopic differences between resources and consumers not only vary between consumer taxa within a functional feeding group but originate from strong and partially contrasting direct effects of riparian vegetation type on resources and consumers (Figure 2,4). To manage dietary imbalances, consumers can use different strategies including compensatory feeding, selective feeding or physiological adaptations (Flores *et al*. 2014; Frainer *et al*. 2016). As many detritivores are feeding on resources of relatively low nutritious value, they likely combine multiple strategies. Food choice experiments have revealed that the degree of compensatory and selective feeding of stream shredders on leaves can vary between taxa (Arsuffi & Suberkropp 1989). Similarly, in our *in situ* stream communities, we find that Crustacea, Plecoptera and Trichoptera shredders have different physiological needs (Frainer et al., 2016) and might therefore use different combinations of strategies to meet them.

Changes at the base of food webs can have cascading effects on higher trophic levels and on biodiversity within and across ecosystems. Studying food-web dynamics beyond ecosystem boundaries will help advance our understanding of the multifaceted effects of land-cover change on biodiversity and ecosystem processes. Further, conservation and restoration practices that account for cross-ecosystem linkages will benefit not only the local biodiversity but also the structure and complexity of food webs and trophic interactions. Protecting and maintaining high biodiversity and intact linkages in ecotones might thus help facing global biodiversity decline and consequences of climate change.

## Acknowledgment

We thank E. Cereghetti, F. Cerotti, L. Sturm, A. R. Esmaeili for their help with fieldwork, and J. Colombo, S. Lötscher, G. Frei, L. Thomas-Sleiman, M. Laurent and S. Hürlemann for assistance in the laboratory. We are also grateful for N. Dubois and I. Brunner for supporting the stable isotope analyses, M. Moretti, P. dos Reis Oliveira and P. Oliveira for their scientific inputs, and all landowners whose property we crossed to access our sampling sites.

## Availability and material

The data that support the findings of this study are openly available on Dryad: LINK

## Ethics statement

All authors confirm that appropriate licenses and permissions have been obtained for invertebrate sampling in Swiss streams.

## Author contribution

Conceptualization: RO, AB, FA; Methodology: RO, AB; Writing: RO, AB, FA; Funding acquisition: AB, FA; Supervision: AB, FA

## Data availability

The data and code that support the findings of this study are openly available on Dryad and Github:

https://datadryad.org/stash/dataset/doi:10.5061/dryad.8pk0p2nv9

https://github.com/RebeccaOester/Riparian-forests-shape-trophic-interactions-in-detrital-stream-food-webs/tree/main

