## Supplementary Material for "Riparian forests shape trophic interactions in detrital stream food webs"

**Running title: Forests shape stream food webs**

**Authors:** Rebecca Oester* [1,2,3], Florian Altermatt [2,3], Andreas Bruder [1]

**Associations**
1 Institute of Microbiology, University of Applied Sciences and Arts of Southern Switzerland, via Flora Ruchat Roncati 15, CH-6850 Mendrisio, Switzerland

2 Department of Evolutionary Biology and Environmental Studies, University of Zurich, Winterthurerstr. 190, CH-8057 Zurich, Switzerland

3 Eawag: Swiss Federal Institute of Aquatic Science and Technology, Department of Aquatic Ecology, Überlandstrasse 133, CH-8600 Dübendorf, Switzerland

Rebecca Oester:

Florian Altermatt:

Andreas Bruder:

### Supplementary Material

Table S1: Hypotheses for links between riparian vegetation type, consumer mass (represented as fungal biomass or shredder body mass) and C:N ratios and $\delta$13C values.

| Link | Hypotheses | Explanation | Source |
|---|---|---|---|
| 1 | For aquatic fungi, we expect **higher** biomass in non-forested sites. | Aquatic fungi may benefit from higher nutrient levels, light, temperatures, and algal growth. | (Danger *et al.* 2013; Paul *et al.* 2006) |
| | For shredders, we expect **lower** average dry weights in non-forested sites. | Shredders may be larger and therefore show higher weight values in more natural/suitable environments. | (Basset *et al.* 2004; Petchey & Belgrano 2010) |
| 2 | For aquatic fungi, we expect **lower** C:N values in non-forested sites. | Aquatic fungi may benefit from higher N levels and temperatures in non-forested streams. | (Brosed *et al.* 2017) |
| | For shredders, we expect **lower** C:N values in non-forested sites. | Shredders may benefit from slightly higher N levels in leaves through increased fungal conditioning in non-forested streams. | (Atkinson *et al.* 2022; Cross *et al.* 2005; Danger *et al.* 2012) |
| 3 | For aquatic fungi, we **expect no difference** in $\delta$13C values between riparian vegetation type. | Aquatic fungi directly incorporate $\delta$13C signatures from the experimental leaves into their biomass and dead leaves likely do not change their $\delta$13C signatures based on environmental conditions. | (Finlay 2001; Goodman *et al.* 2006; Malosso *et al.* 2004) |
| | For shredders, we expect $\delta$13C values to **lie further away** from $\delta$13C signatures of leaves (more positive) in non-forested sites. | Leaves are likely the most important energy source to shredders at forested and non-forested headwater sites, yet in non-forested sites other resources might be important too. | (England & Rosemond 2004; Finlay 2001; Leberfinger *et al.* 2011) |
| 4 | For aquatic fungi, we expect a **negative correlation** between fungal biomass and C:N ratios of leaves. | During decomposition, leaf stoichiometry can vary substantially and with increasing fungal biomass, more N could be accumulated in the leaves. | (Camenzind *et al.* 2021; Manning *et al.* 2015) |
| | For shredders, we expect **no difference** between average dry weight and C:N. | Shredders generally have a very stable C:N ratio due to homeostatic constraints. | (Cross *et al.* 2003; Evans-White *et al.* 2005; Frost *et al.* 2005) |
| 5 | For aquatic fungi, we expect **no difference** between biomass and $\delta$13C values. | Aquatic fungi on the experimentally exposed leaves likely use this substrate as main energy source independent of time and biomass and show no or small carbon isotopic fractionation. | (Costantini *et al.* 2014; Semenina & Tiunov 2010) |
| | For shredders, we expect that $\delta$13C values **depend on consumer mass.** | During the aquatic life stage, shredders typically need various food sources depending on their size and stage-dependent feeding strategies. | (Danger *et al.* 2012; Graça *et al.* 1993) |

Table S2: Detailed information for the seven shredder taxa between forested and non-forested sites with N representing number of measurements (i.e., leaf litter bags with enough biological material).

| Order | Taxon | N | Average abundance per leaf litter bag | | Average dry weight per individual [mg] | |
|---|---|---|---|---|---|---|
| | | | Forested | Non-Forested | Forested | Non-Forested |
| Crustacea | | | | | | |
| | *G. fossarum* | 47 | 14.34±14.24 | 14.29±11.76 | 7.08±7.89 | 10.55±4.63 |
| Plecoptera | | | | | | |
| | *Capnia* sp. | 39 | 19.31±12.51 | 8.00±3.33 | 0.36±0.15 | 0.62±0.08 |
| | *Leuctra* sp. | 14 | 9.50±4.20 | 8.60±2.76 | 0.35±0.07 | 0.37±0.06 |
| | *Nemoura* sp. | 55 | 16.38±8.25 | 18.31±16.04 | 0.32±0.13 | 0.40±0.16 |
| | *Protonemura* sp. | 33 | 142.71±192.75 | 21.47±14.62 | 0.77±0.65 | 1.06±0.30 |
| Trichoptera | | | | | | |
| | Limnephilini | 27 | 2.40±1.58 | 2.59±2.09 | 13.96±18.89 | 14.61±18.34 |
| | Limnephilidae group Auricollis | 71 | 2.42±1.57 | 1.90±1.07 | 6.42±3.54 | 8.97±7.25 |

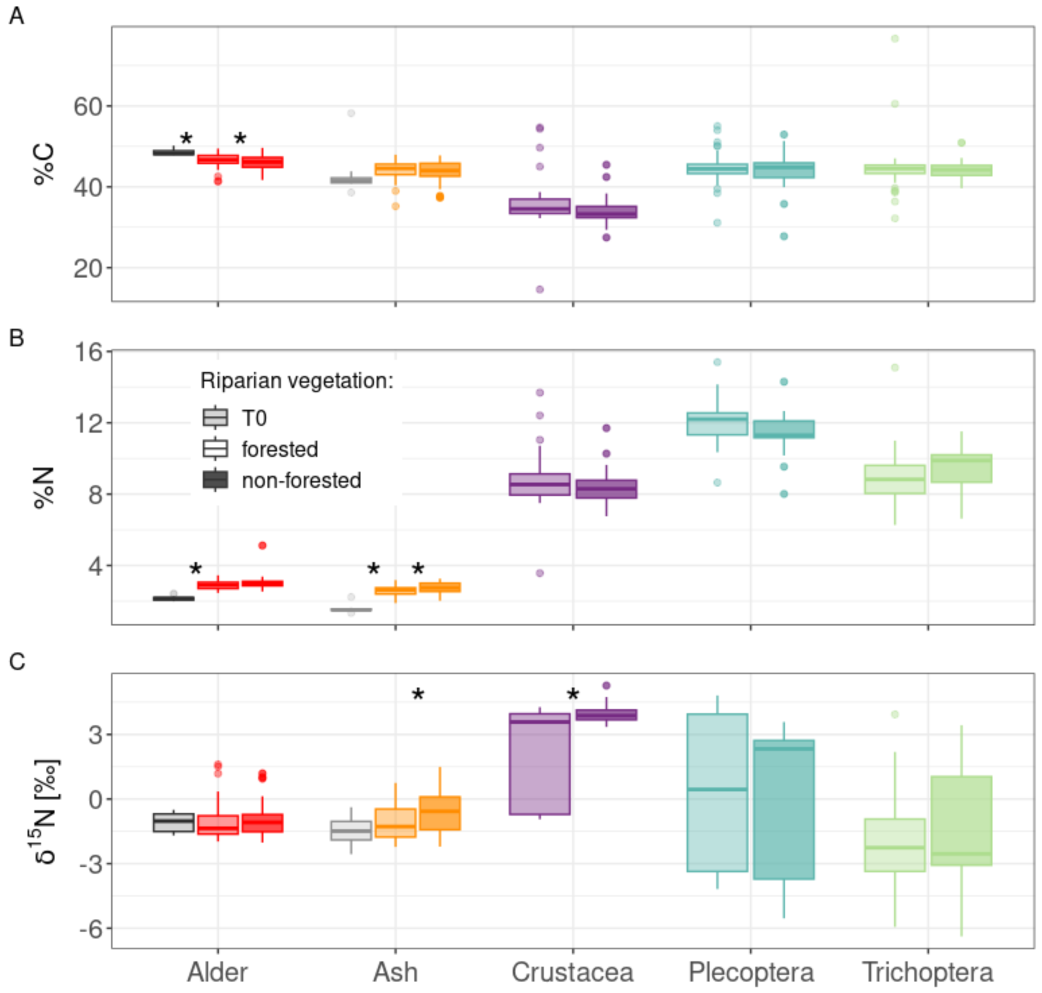


Figure S1: Effects of riparian vegetation and conditioning on resources and consumers A) %C: B) %N and C) $\delta$15N between forested and non-forested sites. Asterisks indicate statistically significant differences between the forested and non-forested sites (i.e., 0 not included in the CIs). Enrichment factors for $\delta$15N values are 2.6 ‰ (Brauns et al. 2018).

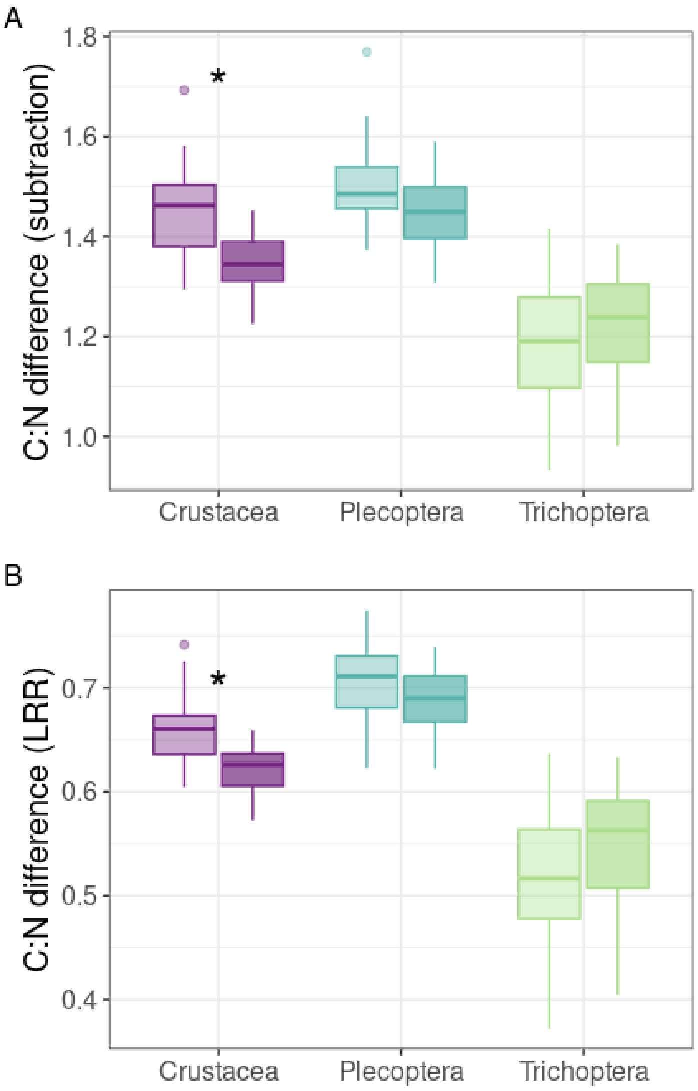


Figure S2: Stoichiometric differences calculated as subtractions (A) and LRR (B). Asterisks indicate statistically significant differences between the forested and non-forested sites (i.e., 0 not included in the CIs).

**Detailed method description for quantifying preservation effects**

To test if the ethanol preservation of macroinvertebrate shredders had an effect on $\delta$13C signatures (Sarakinos *et al.* 2002), we compared $\delta$13C values of preserved and fresh shredders. As a detailed taxonomic determination of many individuals was only possible under a dissecting microscope and under an extended period of time requiring taxonomic expertise, the animals had to be conserved and in 98 % ethanol. We additionally collected 23 macroinvertebrate samples of the same orders from the same sites with kick net sampling in local leaf packs resembling our leaf litter bags and directly dried (60 °C for 48 h) and homogenized the “fresh” individuals.

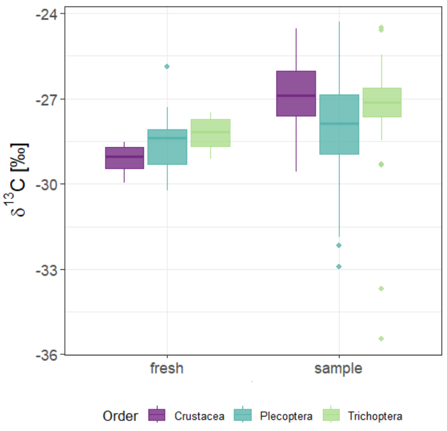


Figure S3: Shift in $\delta$13C values across shredder orders depending on preservation technique with “fresh” indicating a direct processing and “sample” representing ethanol preservation.

We observed a constant shift of 1.09 ‰ of $\delta$13C values and order not being a statistically significant factor (i.e., 0 always included in the CIs). The inclusion or exclusion of this preservation correction factor had no significant effect of the general outcome of our results involving $\delta$13C and interpretations thereof as it just represented a shift of the absolute $\delta$13C values.

Sarakinos, H. C., Johnson, M. L., & Vander Zanden, M. J. (2002). A synthesis of tissue-preservation effects on carbon and nitrogen stable isotope signatures. *Canadian Journal of Zoology* 80:2: 381-387.
